# Agreement between measurements of stance width using motion capture and center of pressure in individuals with and without Parkinson’s disease

**DOI:** 10.1101/122523

**Authors:** J. Lucas McKay

**Affiliations:** The Wallace H. Coulter Department of Biomedical Engineering, Emory University and the Georgia Institute of Technology

**Author notes:** Correspondence to:* J. Lucas McKay, Ph.D., M.S.C.R., Wallace H. Coulter Department of Biomedical Engineering, Emory University and the Georgia Institute of Technology, Room R154, Emory Rehabilitation Hospital, 1441 Clifton Road, Atlanta, Georgia, 30322 USA. Conflict of interest statement:* The author has no conflicts of interest to declare.

**Keywords:** Postural control, Center of pressure location, Measurement, Methodology, Foot position

## Abstract

**Background:** Many individuals with Parkinson’s disease exhibit narrow stance width during balance and gait. Because of this, stance width is an important biomechanical variable in many studies. Measuring stance width accurately using kinematic markers in parkinsonian patients can be problematic due to occlusions by research staff who must closely guard patients to prevent falls.

**Methods:** We investigated whether a measure of stance width based on the mediolateral distance between the center of pressure under each foot could approximate stance width measured with kinematic data. We assessed the agreement between estimates of stance width obtained from simultaneous kinematic and center of pressure measures during quiet standing in 15 individuals (n=9 parkinsonian, n=6 age-similar neurotypical). The source data (1363 unique trials) contained observations of stance width varying between 75–384 mm (≈25-150% of hip width).

**Findings:** Stance width estimates using the two measures were strongly correlated (*r* = 0.98). Center of pressure estimates of stance width were 48 mm wider on average than kinematic measures, and did not vary across study groups (F_2,12_=1.81, *P*<0.21). The expected range of differences between the center of pressure and kinematic methods was 14–83 mm. Agreement increased as stance width increased (*P*<0.02).

**Interpretation:** It is appropriate to define stance width based on center of pressure when it is convenient to do so in studies of individuals with and without Parkinson’s disease. When comparing results across studies with the two methodologies, it is reasonable to assume a bias of 48 mm.

## 1. Introduction

Many individuals with Parkinson’s disease (PD) exhibit narrow stance width during balance and gait (1). Clinically, “narrow stance” is a postural abnormality in which the feet are placed substantially medial to the anterior superior iliac spines (ASIS) (2). Stance width is therefore an important variable in many studies of parkinsonian posture and balance (e.g., (3-5)). It is typically treated as a nominal single value or as a range of values described by the mediolateral distance between kinematic markers placed on the heels, or between the medial malleoli (3-5).

Due to repeated protective steps, dyskinesias, and other practical concerns when studying parkinsonian balance, it is difficult to control stance width precisely during experiments – and so ideally, stance width should be measured as a continuous covariate throughout an experiment. However, doing so with kinematic markers can be problematic due to occlusions by research staff who must carefully guard patients to prevent falls.

Here, we investigated whether a proxy measure of stance width based on the mediolateral distance between the centers of pressure (CoP) beneath each foot could approximate stance width measured kinematically. As typically defined (6), the CoP is the point location of the vertical ground reaction force vector beneath the entire body, and represents a weighted average of all the pressures over the surface area in contact with the ground (6). Whole-body CoP location is often calculated as an important outcome variable in clinical balance studies (5, 7, 8). If bilateral force plates are used, CoP can be calculated separately for each foot (e.g., as it is in instrumented treadmill studies (9)). Since the CoP of each foot must be located within its boundaries, the mediolateral distance between them must be considerably associated with the stance width between the heels during bipedal standing.

We used the approach suggested by Bland and Altman (10) to assess agreement between stance width estimated from foot CoP and measured kinematically in neurotypical individuals (NT) and in parkinsonian individuals in the ON (PD-ON) (8) and OFF (PD-OFF) (11) medication states. We quantified the bias and expected range of differences associated with using stance width estimates from foot CoP rather than kinematic measures. Then, we tested whether differences between methods were associated with group membership (NT vs. PD-ON vs. PD-OFF), and whether differences varied with stance width (12).

## 2. Materials and Methods

### 2.1 PARTICIPANTS

We used baseline measurements from a convenience sample of participants in previous (3) and ongoing cohort studies investigating the effects of rehabilitation on balance responses (Table 1). PD participants were mild-moderate with bilateral symptoms (Hoehn and Yahr stage 2-3 (13)). All participants provided written informed consent and all study procedures were approved by Institutional Review Boards at the Georgia Institute of Technology and Emory University.

**Table 1.** Demographic, clinical, and anthropometric features of the study population.

| Participant | Hoehn & Yahr Stage | Age | Sex | Height, m | Weight, kg | Inter-ASIS distance, cm | Left leg length, cm | Right leg length, cm |
| --- | --- | --- | --- | --- | --- | --- | --- | --- |
| Neurotypical |  |  |  |  |  |  |  |  |
| NT1 | - | 54 | F | 1.62 | 66.7 | 31.3 | 86.5 | 88.5 |
| NT2 | - | 56 | F | 1.78 | 74.8 | 24.7 | 98.0 | 98.0 |
| NT3 | - | 58 | M | 1.64 | 67.2 | 25.5 | 82.5 | 82.0 |
| NT4 | - | 64 | M | 1.80 | 95.2 | 22.0 | 91.0 | 91.8 |
| NT5 | - | 70 | F | 1.57 | 51.0 | 22.3 | 82.0 | 82.0 |
| NT6 | - | 77 | M | 1.85 | 81.9 | 27.8 | 106.0 | 105.0 |
| PD-ON |  |  |  |  |  |  |  |  |
| PDON1 | 2 | 68 | M | 1.80 | 80.9 | 28.5 | 93.5 | 94.0 |
| PDON2 | 2 | 69 | F | 1.55 | 74.8 | 30.1 | 84.0 | 83.0 |
| PDON3 | 3 | 73 | F | 1.80 | 62.7 | 21.1 | 90.0 | 91.0 |
| PDON4 | 2.5 | 79 | M | 1.68 | 68.2 | 27.5 | 91.0 | 90.0 |
| PDON5 | 3 | 79 | M | 1.70 | 74.4 | 24.0 | 89.5 | 90.0 |
| PD-OFF |  |  |  |  |  |  |  |  |
| PDOFF1 | 3 | 75 | F | 1.54 | 50.3 | 22.3 | 83.0 | 82.0 |
| PDOFF2 | 2 | 53 | M | 1.75 | 86.2 | 25.1 | 90.4 | 89.7 |
| PDOFF3 | 2 | 54 | F | 1.63 | 66.0 | 26.2 | 88.0 | 88.0 |
| PDOFF4 | 3 | 82 | F | 1.68 | 59.9 | 25.5 | 94.8 | 94.1 |

Abbreviations: ASIS, anterior superior iliac spine; PD-ON, PD participants in the ON medication state; PD-OFF, PD participants after 12+ hours of withdrawal of antiparkinsonian medications.

### 2.2 EXPERIMENT

As in previous studies (3, 14), participants stood barefoot on two laboratory-grade force plates(AMTI-OR6-6-1000, AMTI, Watertown, MA, USA). The force plates were mounted onto a custom translation platform; however, analyses here considered only periods during which the platform was stationary. Force and moment data were sampled at 1080 Hz and used to calculate the locations of the center of pressure beneath each foot using calibration values supplied with the plates (15-17). Kinematic data were collected at 120 Hz using a Vicon motion capture system (Centennial, CO, USA) and a 25-marker set including reflective markers placed on the left and right heels. Average foot CoP locations and heel marker positions were calculated over the first 250 ms of each trial.

Stance width was controlled by requesting participants press an object (typically a book) between the medial surfaces of their feet, which was subsequently removed before data collection (≈87% of trials), or by manipulating participant’s feet so that kinematic markers on the heels were aligned in the mediolateral direction with tape marks on the floor (≈13%).

### 2.3 DATA ANALYSIS

Stance width measurements derived from CoP and kinematic data were plotted against each other and examined visually. After visual assessment of outliers, trials were excluded due to: 1) tension in a ceiling-mounted fall arrest tether interfering with CoP calculation (17 trials in one participant), and 2) absent video records preventing trial review (2 trials in one participant). After applying exclusions, 1363 trials (41 – 161 per participant) were available for analysis. Stance widths were expressed in mm and normalized to inter-ASIS distance.

Following Bland and Altman (10), correlation between the two measurements was assessed with the Pearson product-moment correlation coefficient *r*. Differences between methods were calculated for each trial and averaged across trials into a single difference value *d_i_* for each participant. Mean values across methods were calculated for each trial and averaged into a single mean value *m_i_* for each participant. Bias between the two methods was quantified as the mean difference *<u>d</u>* (CoP – kinematic method) and the standard deviation of the differences *s*. The limits of agreement were calculated as the range *<u>d</u>*−2*s* to *<u>d</u>*+2*s*. Variation of differences *d_i_* across groups was assessed with one-way ANOVA. Associations between differences *d_i_* and mean values *m_i_* were assessed with *r* (12). Data processing was performed in Matlab (r2016b, The Mathworks, Natick, MA, USA). Statistical procedures were performed in SAS Studio (3.5, The SAS Institute, Cary, NC, USA) and considered significant at *P* = 0.05.

## 3. Results

Stance widths measured from kinematic data varied between 75 – 348 mm, corresponding to 24.9 – 154.1% of inter-ASIS distance. CoP and kinematic stance width measurements are presented in Figure 1A. The two measures were strongly correlated (*r* = 0.98). The mean difference *<u>d</u>* between methods was 48 mm, and the standard deviation of the differences (*s*) was 17 mm. Differences *d_i_* did not vary across groups (F_2,12_=1.81, P<0.21). The limits of agreement, defined as the range *<u>d</u>*−2*s* to *<u>d</u>*+2*s* (10), was 14–83 mm. A “Bland-Altman plot” of the differences between the two methods *d_i_* against their means *m_i_* is presented in Figure 1B. *d_i_* and *m_i_* were significantly negatively correlated (*r* = −0.59, P<0.02).

**Figure 1.**
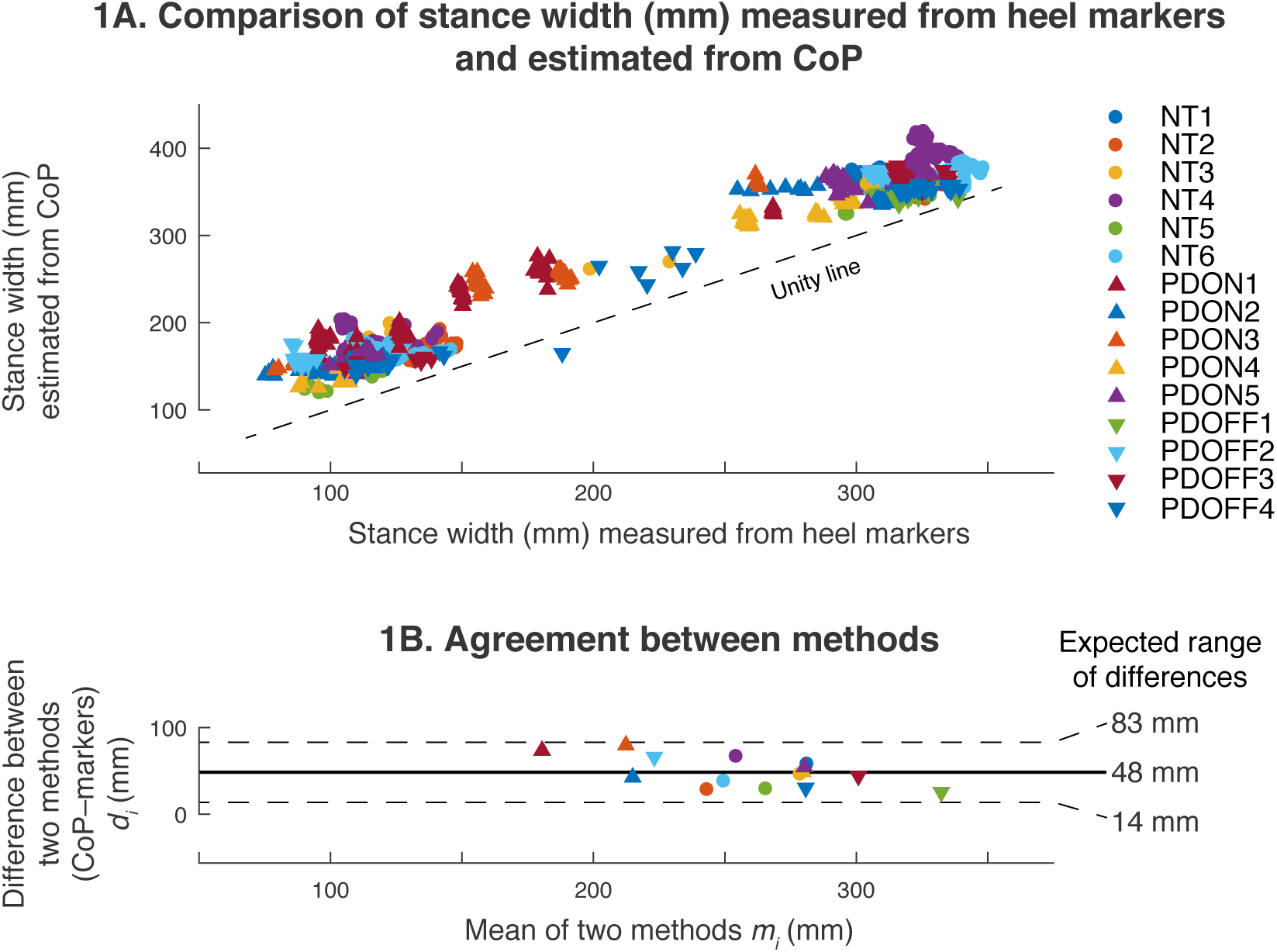
Comparison of stance width measurements from kinematic and CoP data. A: Plot of results of one method (CoP, ordinate) against those of the other (kinematics, abscissa). Marker shapes designate study group and participants are coded by color. B: “Bland-Altman” (10) plot of limits of agreement between the two methods. The CoP method introduces an absolute bias *<u>d</u>* of 48 mm and an expected range of deviations 14-83 mm. Color and marker codes are as in part A.

## 4. Discussion

Stance width is an important variable in many studies of parkinsonian (4, 5) and neurotypical (18, 19) posture and balance. We found that stance width estimates from foot CoP and kinematic markers were strongly linearly correlated, and that on average, measures of stance width derived from CoP were 48 mm wider than those derived from kinematic markers. This bias that can be explained by the externally-rotated “toe out” posture used by most participants, in which a substantial portion of the foot plantar surface lies lateral to the posterior face of the heel. Overall, these results suggest that foot CoP location, a commonly calculated variable in clinical biomechanics studies (5, 7, 8) can be used to approximate stance width in healthy aging and in individuals with PD in the ON and OFF medication states.

We noted that differences between methods were non-negligible – ranging from 14 to 83 mm. However, this precision is adequate to discriminate between nominal stance widths used in the literature, which are typically separated by 100 mm or more (4, 18). Due to the high precision of CoP calculation with laboratory force plates (2-5 mm (17)), the primary source of variability in differences is probably trial-to-trial variability in weight distribution, rather than instrumentation error.

There are two notable limitations to this approach. First, differences between methods were highest at the narrow stance widths preferred by PD subjects, a fact that should be considered carefully during study design. Second, because these participants were allowed to adopt a comfortable “toe out” orientation during testing, the agreement between the methods in experimental paradigms in which foot orientation is enforced (e.g., parallel (4); 20° (18)) remains to be established.

## 5. Conclusion

In summary, these results suggest that: 1) it is appropriate in studies of individuals with and without PD to define stance width based on CoP, and 2) when comparing results across studies with the two methods, it is reasonable to assume a bias of 48 mm.

## Acknowledgements

I thank Dr. Jessica Allen, Dr. Jeffrey Bingham, Dr. Madeleine Hackney, Dr. Lena Ting, and Ms. Kimberly Lang for their expertise and assistance.

